# The model of local axon homeostasis - explaining the role and regulation of microtubule bundles in axon maintenance and pathology

**DOI:** 10.1101/577320

**Authors:** Ines Hahn, André Voelzmann, Yu-Ting Liew, Beatriz Costa-Gomes, Andreas Prokop

**Affiliations:** The University of Manchester, Manchester Academic Health Science Centre, Faculty of Biology, Medicine and Health, School of Biology, Manchester, UK

**Keywords:** *Drosophila*, neurodegeneration, axons, actin, cytoskeleton, microtubules

## Abstract

Axons are the slender, cable-like, up to meter-long projections of neurons that electrically wire our brain and body. In spite of their challenging morphology, they usually need to be maintained for an organism’s lifetime. This makes them key lesion sites in pathological processes of ageing, injury and neurodegeneration. The morphology and physiology of axons crucially depends on the parallel bundles of microtubules (MTs), running all along to form their structural backbones and highways for life-sustaining cargo transport and organelle dynamics. Understanding how these bundles are formed and then maintained will provide important explanations for axon biology and pathology. Currently, much is known about MTs and the proteins that bind and regulate them, but very little about how they functionally integrate to regulate axons. As an attempt to bridge this important knowledge gap, we explain here the model of local axon homeostasis, based on our own experiments and published data. (1) As the default, we observe that axonal MTs have a strong bias to become disorganised, likely caused by the physical forces imposed by motor proteins and their life-sustaining functions during intra-axonal transport and dynamics. (2) Preventing MT disorganisation and promoting their bundled conformation, requires complex machinery involving most or even all major classes of MT-binding and - regulating proteins. As will be discussed, this model offers new explanations for axonopathies, in particular those linking to MT-regulating proteins and motors; it will hopefully motivate more researchers to study MTs, and help to decipher the complex regulatory networks that can explain axon biology and pathology.

## Introduction

Axons are the slender, cable-like extensions of nerve cells which form the nerves and nerve tracts that wire our brain and body, sending neuronal messages in highly regulated manners. With diameters of only 0.1-15μm (Hoffman, 1995), they extend over distances of up to a meter in humans. To adopt such a unique morphology and physiology, axons display many specialised features (Fig.1).

**Fig. 1.**
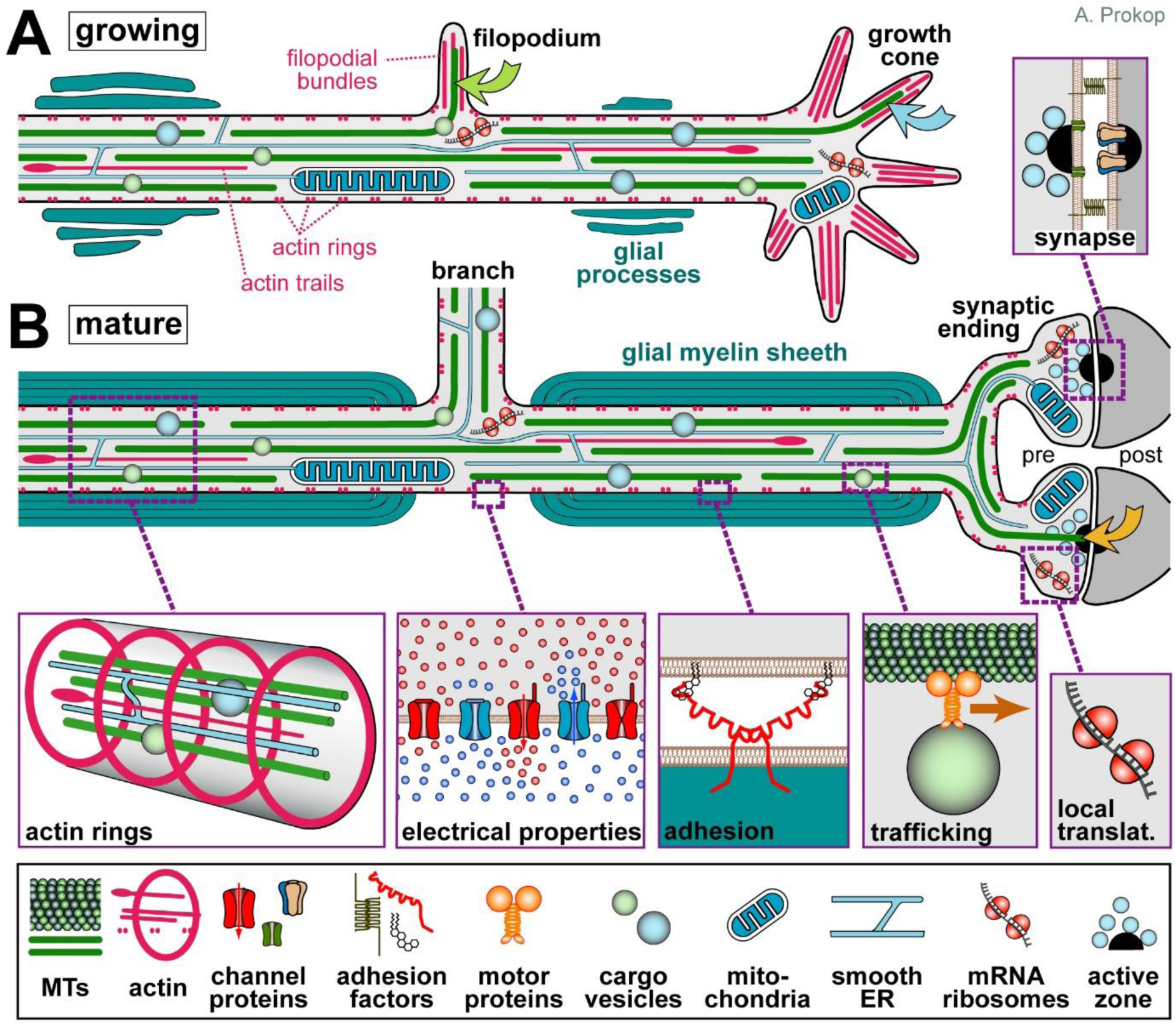
Specific properties of axons. Axons during the growth cone stage are shown in (**A**) and after synaptic maturation in (**B**), differing primarily in certain stage-specific specialisations including growth cones, synapses, electrical properties and glial interactions (here myelination; Meyer and Kaspar, 2017; Pan and Chan, 2017). The core machinery in the axon shaft can be expected to be similar at both stages: parallel continuous bundles of extended but discontinuous MTs run all along axons serving as a structural backbone (see Fig.2), a transport highway for axonal trafficking (driven by motor proteins), and a source for ‘off-track’ MTs contributing to morphogenetic processes including branch formation, directed axon growth and synapse formation/plasticity (green, orange, blue curved arrows); MT bundles are interspersed with longitudinal actin trails (Leterrier et al., 2017), continuous networks of (smooth) ER (Gonzalez and Couve, 2014), and other membranous organelles including mitochondria (Saxton and Hollenbeck, 2012); axonal membranes display regularly spaced periodic rings of cortical actin (Qu et al., 2017; Xu et al., 2013; Vassilopoulos et al., 2019), an unusually high number of ion-specific channel proteins and transporters to conduct nerve impulses (Kandel et al., 2012), as well as adhesions with external structures including parallel axons (not shown), glial processes (Pronker et al., 2016) and synaptic partner cells (Koper et al., 2012); a degree of independence from cell-body derived proteins is provided by local translation machinery (Cioni et al., 2018; Giuditta et al., 2002b; Shigeoka et al., 2018) or supply from surrounding glia cells (not shown; Court et al., 2011; Frühbeis et al., 2013; Giuditta et al., 2002a; Rajendran et al., 2014). Note that the axon diameter in the region between glia cells in B (referred to as Node of Ranvier) usually has a much smaller diameter than the rest of the axon (Hoffman, 1995).

Axons are indispensable for nervous system function, as illustrated by paralysis in spinal cord injury caused by the interruption of ascending and descending axon tracts (Bichenback, 2013; Tedeschi and Bradke, 2016). Axons are key lesion sites in injury-induced trauma and coma (Gaetz, 2004; Medana and Esiri, 2003; Smith et al., 2000; Tang-Schomer et al., 2012), and axon decay is believed to be an important trigger for neuronal loss in ageing and many neurodegenerative disorders (Adalbert and Coleman, 2012; Salvadores et al., 2017). Notably, most neurons cannot be replaced, and compensation of lost axons through collateral branching of intact neighbouring axons has obvious limitations (Adalbert and Coleman, 2012; Sturrock, 1987).

This means that most axons have to be maintained for an organism’s life time, i.e. up to a century in humans; unsurprisingly, mammals tend to lose almost half their axon mass towards high age (Calkins, 2013; Marner et al., 2003). This trend is severely enhanced in neurodegenerative disorders, as illustrated by gradually increasing paralysis in spastic paraplegia or motorneuron disease (Blackstone et al., 2011; Riancho et al., 2019).

Research into neurodegenerative disorders typically approaches the problem by describing observed phenotypes and unravelling the molecular mechanisms performed by proteins linked to the disease. However, this approach rarely leads to satisfactory explanations of the pathology (Aguzzi, 2019). We believe that more profound understanding will arise when widening the scope from molecular to cellular mechanisms, by studying how proteins work within regulatory networks to underpin observable processes of axon biology - thus performing investigations at the same level of complexity at which pathology becomes manifest. Here we will illustrate this approach by focussing on the axonal cytoskeleton.

### The importance of microtubule bundles for axon biology

As illustrated in Fig. 1, the cytoskeleton of the axon shaft consists of straight parallel bundles of MTs, which are interspersed with intermediate filaments (not shown) and longitudinal actin fibres called ‘actin trails’ - all running through evenly spaced periodic rings, proposed to consist either of short and adducin-capped actin filaments (Qu et al., 2017; Xu et al., 2013) or of two long intertwined actin filaments (Vassilopoulos et al., 2019) - future will show. Significant deviations from this organisation that will not be considered in this review, exist at axon initial segments (not shown in Fig.1), growth cones and synapses (Dent et al., 2011; Leterrier, 2018; Leterrier et al., 2017; Prokop, 2013).

Of the three cytoskeleton classes, intermediate filaments were suggested by anatomical, developmental and genetic studies to regulate axon diameter, and their axonal aggregation is a hallmark of many neurodegenerative diseases (Friede and Samorajski, 1970; Hoffman, 1995; Perrot et al., 2008; Rao et al., 2003; Sakaguchi et al., 1993). However, intermediate filament accumulations are not necessarily the cause, but can be the consequence of axon decay (Eyer et al., 1998; Nguyen et al., 2000; Perrot et al., 2008). Notably, *Neurofilament-H-lacZ* mutant mice or *Quiver* mutant quail completely lack axonal intermediate filaments, but develop and breed fairly normally (Eyer and Peterson, 1994; Yamasaki et al., 1991). Furthermore, various arthropods form axons of defined diameters in the absence of any axonal intermediate filaments (Allen et al., 2006; Hirokawa, 1986; Voelzmann et al., 2016a). In contrast to the moderate roles of intermediate filaments, actin and microtubules (MT) are essential for all stages of neuronal development and maintenance (Sakakibara et al., 2013; Tas and Kapitein, 2018; Voelzmann et al., 2016a); this review will be dedicated to the role and regulation of MTs.

Axons contain bundles of MTs that run along the entire length of their shafts (Fig.1); these bundles are essential for axon biology in at least three ways: as structural backbones, as highways for axonal transport and organelle dynamics, and as source for splaying MTs that can contribute to axon morphogenesis or physiology (details in Box 1). Maintaining MT bundles is therefore crucial for axon longevity. Accordingly, there are prominent and numerous genetic links from MT regulators to hereditary neurodegenerative disorders (Suppl. Mat. in Prokop et al., 2013), and axon decay is a frequent side effect of MT-targeting chemotherapies (Prior et al., 2017; Wozniak et al., 2018; Wu et al., 2014). Of particular interest for this review are reports of pathological axon swellings where MT bundles have disintegrated into loops or waves (bottom of Fig.3), occurring in ageing, after injury and in certain axonopathies (Adalbert et al., 2009; Bernier and Kothary, 1998; Dalpe et al., 1998; Denton et al., 2014; Fassier et al., 2013; Havlicek et al., 2014; Sorbara et al., 2014; Tang-Schomer et al., 2012; Tarrade et al., 2006; Yamasaki et al., 1991; Yin et al., 2016). Notably, one study suggests that MT aberration upon ageing could cause swellings that trap and damage mitochondria, thus triggering axon degeneration (Fiala et al., 2007). However, in the existing literature too little emphasis is given to MTs and there are simply not enough data to deduce meaningful correlations between axon degeneration and MT bundle decay.

**Fig. 2.**
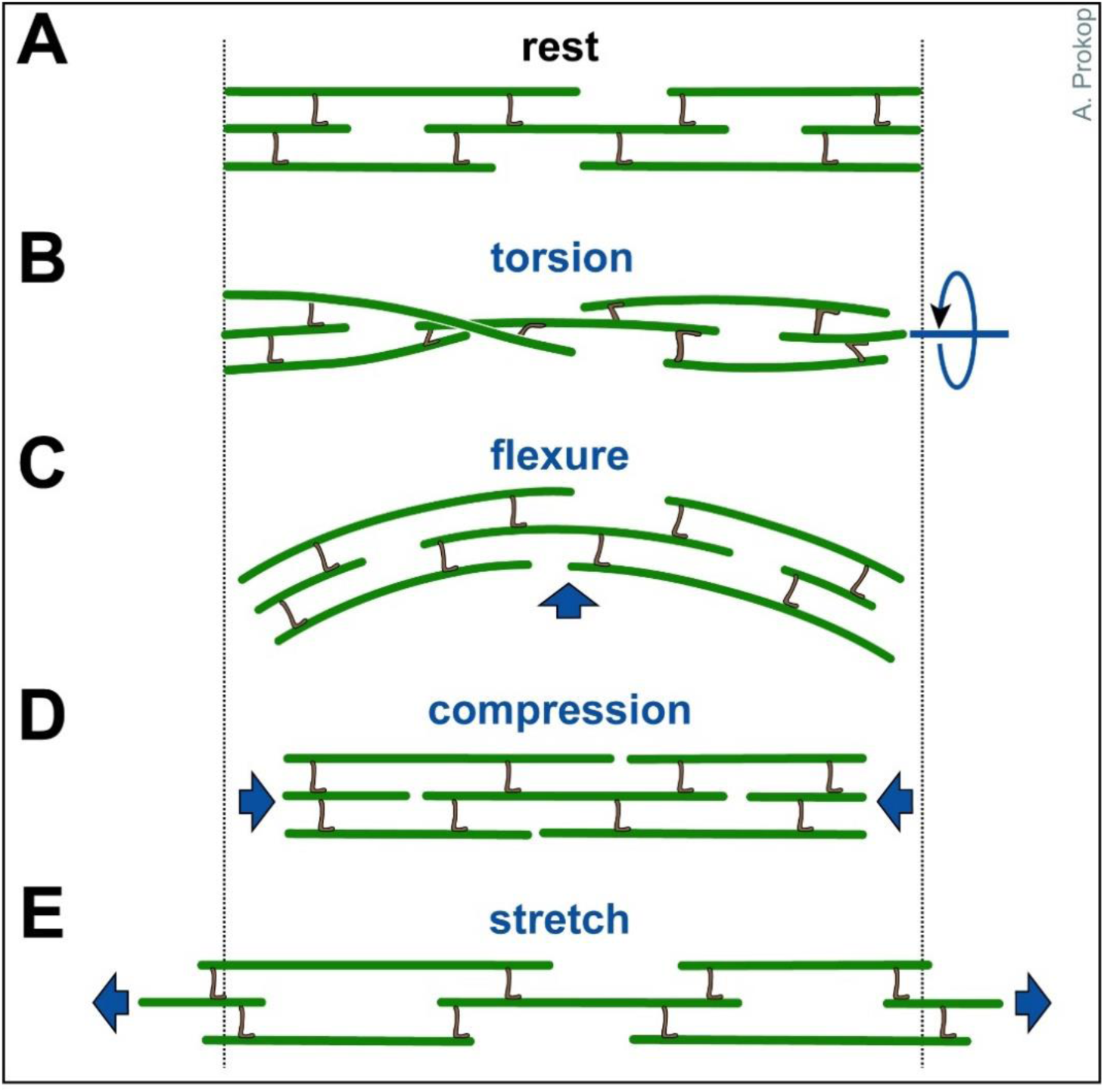
Axonal response to mechanical challenges. Continuous bundles of discontinuous MTs which are flexibly cross-linked (likely involving slip-bonds) are thought to provide a structural element that can respond to different forms of mechanical impact (as indicated in blue).

**Fig. 3.**
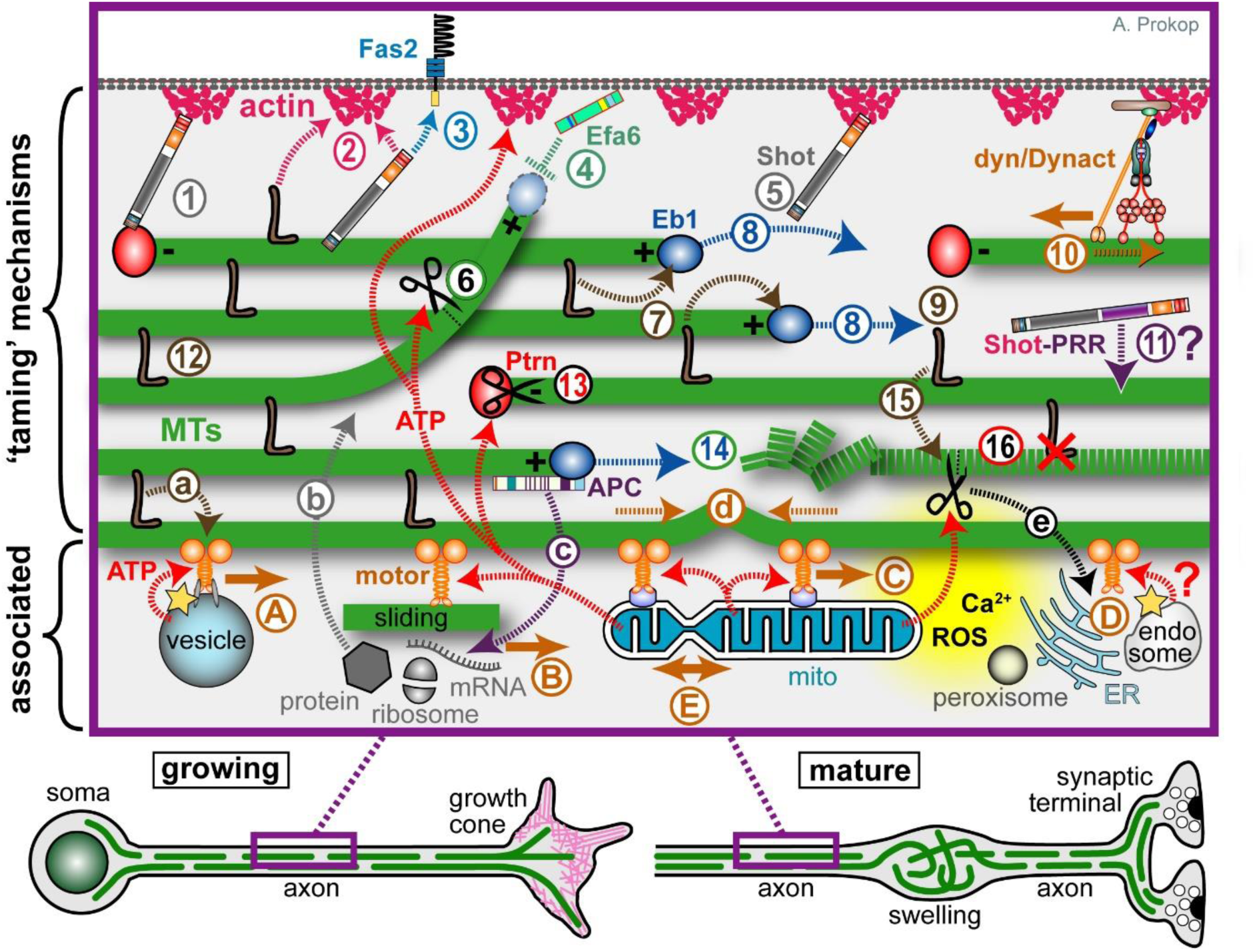
An interactome of MT-regulating and-associated mechanisms expected to contribute within the model of local axon homeostasis. Developing and mature neurons are shown at the bottom indicating that the close-up (magenta frame) might apply in both contexts. **1**-**16**) Potential mechanisms that can ‘tame’ MTs into bundled conformation: MT polymerisation (blue stippled arrows) is driven by molecular machinery centred on Eb1 (blue balls), further influenced by the tubulin-supply machinery (not shown) and shaft-binding proteins (**7**); polymerisation generates new MTs required for bundle formation (**8**) and turn-over (**14**); to integrate into bundles, extending MTs require guidance via actin-Eb1 cross-linkage along the axonal surface (**5**; Shot) or along pre-existing MTs through MT-MT cross-linkers (**9**; brown L). The same or other cross-linkers provide the structural glue that holds MT bundles together (**12**; brown L); some of them can also bind to actin (**2**), they protect from (or recruit) MT severing activity (**15**), and influence motor protein dynamics (**a**). MTs which have escaped any cross-linkage are eliminated by cortical collapse factors when approaching the axonal surface (**4**; Efa6) or by MT severing factors at MT-MT cross-points (**6**). The bundled MTs are discontinuous; their free minus ends are stabilised by CAMSAP/Patronin (Ptrn) together with katanin (black scissors; **13**), whereas non-polymerising MT plus ends are stabilised by other factors (not shown; e.g. CLASP or the Dynactin subunit p150/Glued; Hur et al., 2011; Lazarus et al., 2013). The dynein/Dynactin complex is believed to link cortical actin to MT bundles and drive them anterogradely (**10**), whereas Ptrn at minus ends may anchor MTs via spectraplakins to the axon cortex (**1**); spectraplakins may also link MTs directly to cortical actin (**2**) or to transmembrane receptors (**3**), and they are expected to perform further, still unexplored actin-independent bundle-promoting roles through their PRR domains (**11**). Tear-and-wear damages MTs (dashed green line), potentially affecting interaction with MT-binding proteins (**16**; red X); MT severing proteins might selectively eliminate such MTs (**16**; scissors) or MTs undergo repair (not shown). **A**-**E**) Mechanisms closely ‘associated’ with MT bundles: MT-associated motor proteins (’motor’, solid orange arrows) drive axonal transport of (protein-loaded) vesicles (**A**), cytoplasmic factors including proteins, translational machinery (ribosomes) or RNAs (**B**), move other MTs (**B**, sliding), and position/rearrange organelles including mitochondria (**C**, mitos), ER, peroxisomes and endosome (**D**) - and this likely includes mitochondrial fission and fusion (**E**). **a**-**e**) The motor-associated functions all act downstream of MT bundles because they require them to walk on; but they also act upstream: for example, the forces they generate (stippled orange arrows) are the potential cause for MT disorganisation (buckling shown in **d**); transport delivers required regulators and building blocks for bundle-maintaining processes (**b**); the proper regulation of organelles/endocytic compartments provides systemic factors that can orchestrate taming mechanisms, including intracellular free calcium or reactive oxygen species (Ca^2+^, ROS; yellow cloud) as well as ATP required for many processes including actin dynamics, MT severing and MT motor activity (red stippled arrows; note that vesicular transport uses glycolysis to generate its own ATP; yellow star); *vice versa*, the MT severer spastin also regulates the ER through ATP-independent mechanisms (**e**), and MT-associated proteins (APC) regulate local translation events (**c**).

#### Box 1 The roles of axonal MT bundles

(1) Axonal MT bundles serve as structural backbones, comparable to the vertebral column of a snake; since MTs in these bundles are discontinuous and expected to be interlinked via flexible connections (see section on cross-linkers), they are ideally suited to respond to longitudinal stretch and compression (similar to a half-extended telescope ladder), but also to torsion and flexure (Fig.2).

(2) Axonal MT bundles provide the highways for life-sustaining axonal transport between cell bodies and the axonal compartment. This transport is driven anterogradely by kinesins and retrogradely by the dynein/Dynactin complex; the cargoes include mRNAs, cytoplasmic proteins including signalling factors, vesicles delivering synaptic proteins, transmembrane proteins, neuropeptides and/or membrane lipids, as well as entire organelles including mitochondria (Fig. 3A-D; Goldstein et al., 2008; Gondre-Lewis et al., 2012; Gonzalez and Couve, 2014; Hirokawa et al., 2010; Pfenninger, 2009). Furthermore, local dynamics of organelles, such as fission or fusion of mitochondria, can be expected to require forces generated by MT-associated motor proteins (Fig. 3E; Saxton and Hollenbeck, 2012).

(3) Axonal MT bundles provide a source for readily available MTs that can be used for other purposes (curved arrows in Fig.1); for example, splaying MTs can trigger axon extension processes in growth cones (Dent et al., 2011; Prokop et al., 2013; Miller and Suter, 2018), induce branching through growth cone splitting (Acebes and Ferrus, 2000) or collateral branch formation along the axon shaft (Kalil and Dent, 2014; Tint et al., 2009; Tymanskyj et al., 2017), as well as support physiological changes at synapses (Bodaleo and Gonzalez-Billault, 2016).

Even if there were a close correlation, this still does not exclude that, depending on the pathological condition, MT bundle deterioration may be a mere consequence rather than cause of axon decay (details in Fig.4). Ultimate clarification will only arise from developing a better understanding of MT bundle-forming and-maintaining machinery. Here we propose a conceptual framework that may facilitate such developments.

**Fig. 4.**
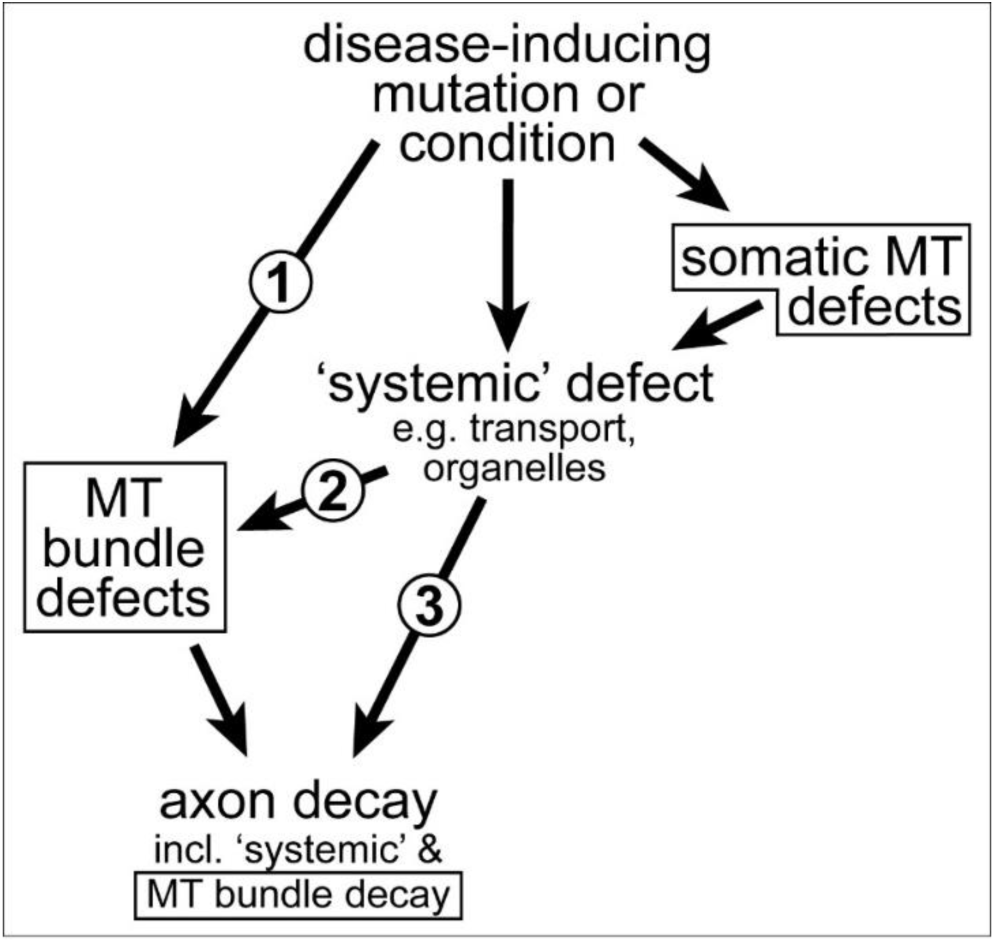
MT bundle defects as cause or consequence of axon decay. **1**) Disease-inducing mutations/conditions can affect a MT-bundle regulator (e.g. dystonin; Voelzmann et al., 2017), thus causing MT bundle defects first which can, in turn, trigger axon decay. **2**) Disease-inducing mutations/conditions can affect systemic factors which, in turn cause MT bundle defects as an intermediate causative step in the cascade leading to axon decay (e.g. axonal transport fails, leading to MT bundle defects which then contribute to axon decay (e.g. Alzheimer’s disease or ALS; Brandt and Bakota, 2017; Farah et al., 2003; Zempel and Mandelkow, 2015); this may occur even if MT regulators are affected, but these regulators mainly act in the cell body (e.g. dysregulation of the Golgi; Ferrier et al., 2013). **3**) MT bundle deterioration may be a mere consequence of axon decay, although this case will be difficult to disentangle from option 2, since MT bundle disintegration and axonal disassembly may occur in parallel, as observed in developmental or injury-induced axon degeneration; Bradke et al., 2012; Wang et al., 2012; Yaron and Schuldiner, 2016). All MT-related phenotypes in this graph are emphasised with a frame.

### From work in *Drosophila* to the integrated model of local axon homeostasis

The foundations for this conceptual framework were laid when we took the decision to use the fruit fly *Drosophila melanogaster* as a means to study how cytoskeletal regulators collaborate in orchestrating the morphogenetic changes that drive axon growth (Sánchez-Soriano et al., 2007). *Drosophila* is not a miniature human, but it has many advantages and provides powerful means to uncover the regulatory concepts behind the roles and regulations of axonal MTs, which then often apply to higher organisms (Box 2; Aguzzi, 2019; Bellen et al., 2010; Elden et al., 2010; Prokop, 2018). Through using *Drosophila* neurons as a consistent standardised cell system, our group alone performed functional analyses of over 50 actin-and/or MT-binding or - associating regulators (Prokop et al., 2013); these studies form an unprecedented pool of data on the basis of which to develop novel concepts (Alves-Silva et al., 2012; Beaven et al., 2015; Gonçalves-Pimentel et al., 2011; Qu et al., 2018; Qu et al., 2017; Voelzmann et al., 2016b).

#### Box 2 Why use *Drosophila*?

The use of *Drosophila* neurons to study the neuronal cytoskeleton has a number of advantages that were detailed elsewhere (Prokop et al., 2013). Key aspects are the high degree of conservation of cytoskeletal proteins, regulators and dynamics, the experimental amenability of neurons in primary cell culture and *in vivo* (Prokop et al., 2013; Prokop et al., 2012; Sánchez-Soriano et al., 2010), and the relative ease of genetic manipulation based on available resources and efficient combinatorial genetics (Hahn et al., 2016). The power of combinatorial genetics is rooted in the relative ease, speed and cost effectiveness with which genes can be manipulated and functionally analysed, facilitating also combined analyses of multiple factors in the same animals or cells (Prokop, 2018; Prokop et al., 2013; Roote and Prokop, 2013). Combinatorial genetics has been extremely successful in overcoming problems of redundancy, and generating new conceptual understanding of co-operative networks of MT regulation (see main text). This can hardly be achieved through isolated work on individual factors.

Our loss-of-function analyses of 24 MT-binding or -associating (2^nd^ order) proteins, revealed that more than half displayed significant MT disorganisation. Interestingly, the MT disorganisation found in these various conditions appears to display certain common characteristics: axons display areas in which their bundles are dissolved into chaotic, intertwined, crisscrossing arrangements of curled MTs (see examples in Fig.5). These phenotypes were surprising when considering that MTs usually behave like rigid rods (Fletcher and Mullins, 2010; Hawkins et al., 2010; Howard, 2001). Notably, when using some of the same genetic conditions *in vivo*, comparable phenotypes were observed in the fly brain (Qu et al., 2018). Such *in vivo* phenotypes in the fly remind of the curled MT conformations in pathological axon swellings of mammalian models mentioned in the previous section. Potential evolutionary conservation of this phenomenon is further supported by the occurrence of similar MT curling and disorganisation in mouse and rat primary neurons (Ahmad et al., 2006; Sánchez-Soriano et al., 2009) - and more reports will emerge once researchers consider MT disorganisation a phenotype worth quantifying, not just an artefact.

**Fig. 5.**
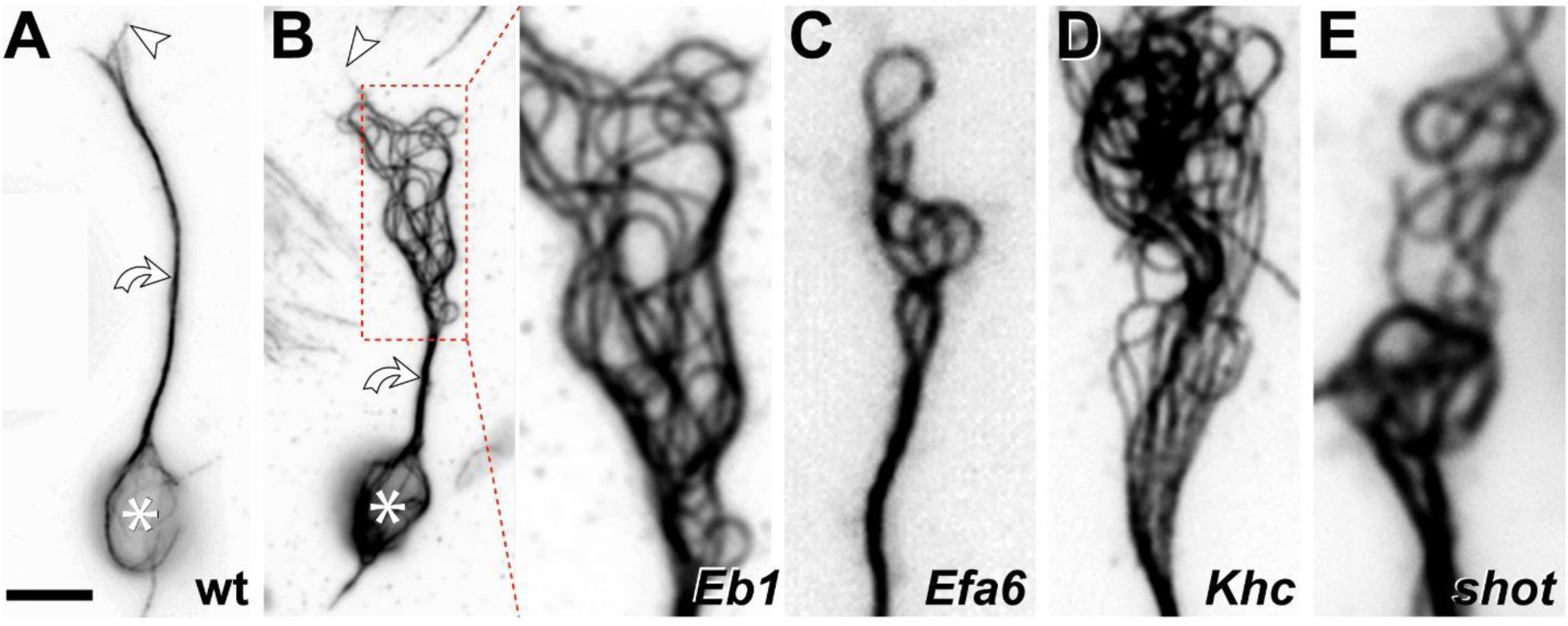
Disorganisation of axonal MTs upon loss of different MT regulators in *Drosophila* primary neurons. **A)** Normal neuron (wild-type, wt) with soma (asterisk), axon shaft (curved arrow) and growth cone (tip of most distal MT indicated by arrow head). **B)** *Eb1^5^* mutant neuron where the area of MT disorganisation is framed by a red stippled box and shown as close-up on the right. **C-E)** Similar close-ups shown for *Efa6^GX6[w-]^*, *Khc^27^* and *shot^3^* mutant neurons. Note that the four mutated factors perform fundamentally different molecular functions, with Eb1 being a MT plus-end binder (’8’ in Fig.3), Efa6 a cortical collapse factor (’4’ in Fig.3), Khc a kinesin-1 motor protein (’A-E’ in Fig.3) and Shot a multi-functional cross-linker (’1-3, 5, 11’ in Fig.3). All neurons were derived from wild-type or homozygous mutant embryos, mechanically and chemically dissociated, kept for 7days in pre-culture in a centrifuge tube to deplete any maternal gene product, mechanically and chemically dissociated again, cultured on concanavalin A-coated glass coverslips for 1day at 21°C, fixed and stained with anti-α-tubulin (DM1A, Sigma; procedures detailed elsewhere: Prokop et al., 2012); images were taken using STED (stimulated emission depletion) microscopy. Scale bar in A represents 10 μm for the two neurons and 4 μm in close-ups.

As an attempt to explain the occurrence of this unusual phenotype across mutant conditions and animal groups, we developed the model of ‘*local axon homeostasis*’ (Prokop, 2016; Voelzmann et al., 2016a), based on two fundamental elements:

1. The model proposes that MTs in axons show a strong bias to become disorganised, most likely because they are challenged to buckle (’d’ in Fig.3) and/or curl up by the narrow axonal environment enriched in MTs, force generating motor proteins and physical obstacles posed by organelles and protein complexes (Fig.1, ‘A-E’ in Fig.3). Once MT disorganisation occurs, it can develop into pathological axon swellings.
2. The model proposes that this risk is contained through the actions of different classes of MT-associating and -regulating proteins, which co-operate and complement each other to form robust machinery that ‘tames’ MTs into bundles (’1-16’ in Fig.3).

In this model, each axon segment uses local action of MT regulators to maintain its bundled MT organisation (hence ‘local axon homeostasis’). Hereditary or acquired loss of single regulators would be expected to weaken this machinery and increase the statistical risk of MT disorganisation. Such heightened probability might explain why many axonopathies affect primarily long axons (Prior et al., 2017), and why certain disorders linked to MT regulators display late onset of axon decay (Voelzmann et al., 2017).

In the next two sections, we discuss potential causes explaining the bias of axonal MTs to become disorganised. We will then summarise experimentally demonstrated MT bundle-maintaining mechanisms, and speculate about further mechanisms based on existing knowledge of known classes of axonal MT-regulating proteins.

### Understanding the unusual curling behaviours of MTs in axons

Although curvature is a key driver of MT plus end dynamics during de-/polymerisation (Brouhard and Rice, 2018; van Haren and Wittmann, 2019), MT lattices *in vitro* usually behave as rigid rods with a persistence length of 1-10 mm (as compared to ∼12 μm measured for actin filaments; Fletcher and Mullins, 2010; Hawkins et al., 2010; Howard, 2001). MTs are polar polymers composed of α/ß-tubulin heterodimers which are arranged in a head-to-tail fashion into linear protofilaments; usually 13 of these protofilaments are laterally aligned forming a straight tube of roughly 25 nm diameter (Fig.6A, C). But MTs can deviate from this norm, and this may introduce an intrinsic element of disorder into MT bundles: for example, axonal MTs were reported to contain 13 protofilaments in frog olfactory or goldfish brain axons, but 11 or 15 in *C. elegans*, and 12 in *Drosophila*, crayfish and lobster (Benshalom and Reese, 1985; Burton et al., 1975; Savage et al., 1989). Deviation from the straight 13 protofilament conformation appears to equip MTs with distinct, functionally relevant physical properties (Chaaban and Brouhard, 2017; Chalfie and Thomson, 1982). But it also introduces a skew into the MT structure, which causes a supertwist of the tubule (Fig.6D; Chrétien and Fuller, 2000; Chrétien et al., 1996; Chrétien and Wade, 1991); this supertwist forces motor proteins to rotate around MTs (Ray et al., 1993) and is the likely explanation for supercoil of entire axons observed upon MT bundle destabilisation (Krieg et al., 2017; Shaw and Bray, 1977).

**Fig. 6.**
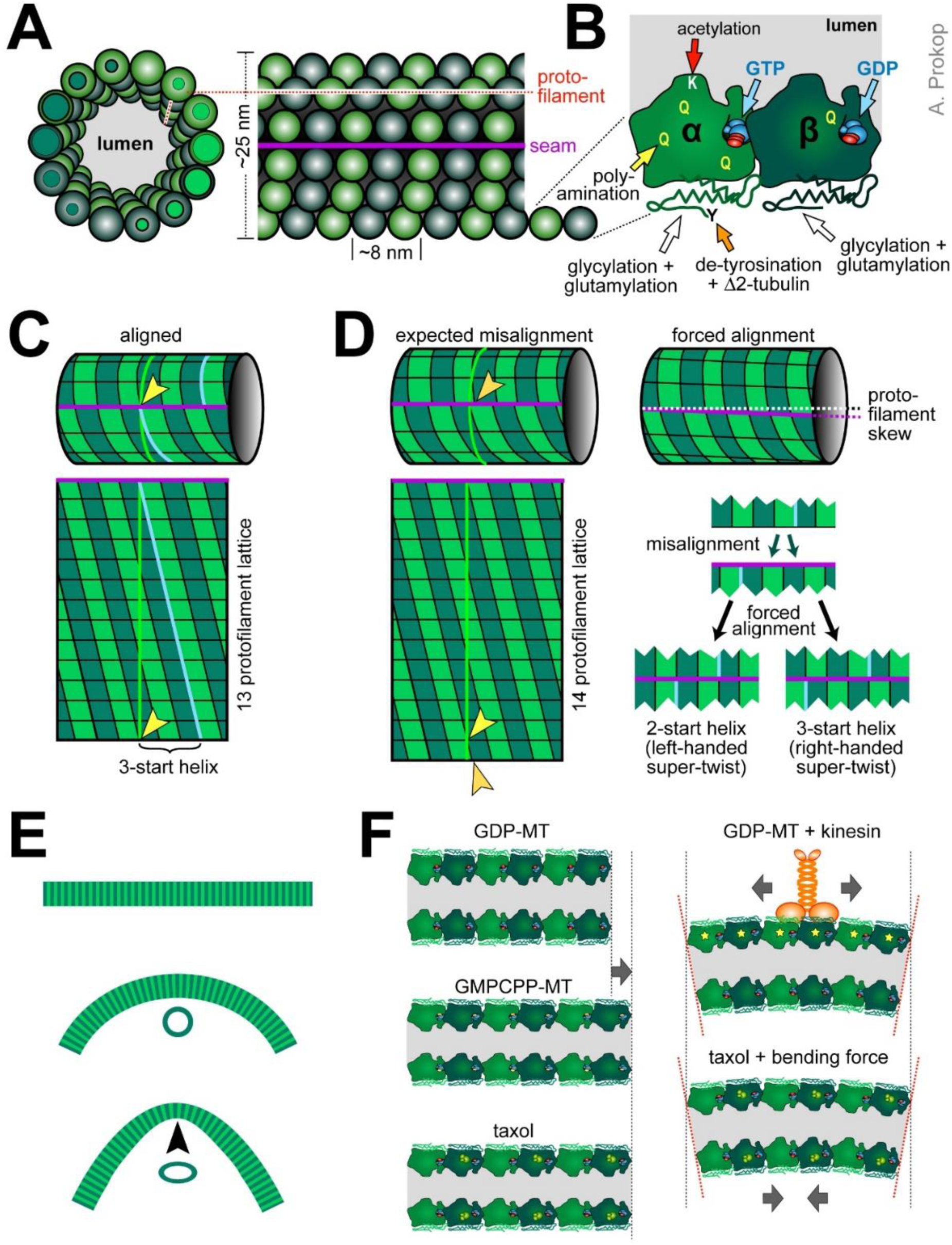
A molecular perspective of microtubule properties. **A)** Cross-section of a MT with 14 protofilaments (PF) and lateral view of a 13 PF MT, both in B-lattice configuration, where α-tubulins make lateral bonds with α-tubulins and ß with ß, except at the seam (magenta line: seam; dashed red line: PF). **B)** Close-up of an α/ß-tubulin heterodimer showing the various post-translational modification sites as indicated; note that the GTP of ß-tubulin in lattices is usually hydrolysed (GDP). **C)** A 13 PF MT (top), cut open at the seam and rolled out (bottom); the yellow line shows the diameter, the blue line follows the helical rise of laterally bonded tubulins; in 13 PF MTs, tubulins are precisely aligned at the seam (yellow arrow head) but shifted by three positions (3-start helix). **D)** When deviating from the 13 PF prototype, tubulins are misaligned at the seam (orange arrow head); when forced into alignment, the PFs skew, causing a super-twist of the MT as described by the ‘lattice accommodation model’ (Chrétien and Fuller, 2000; Langford, 1980); for certain PF numbers, MTs can form two alternative alignments, of which usually the version with the lower helix start value (left) has a left-handed super-twist, the higher value is right-handed (Chrétien and Fuller, 2000). **E)** MTs behave like rigid rods with a persistence length of up to 10 mm, but can be bent down to diameters of ∼1μm before they break; it has been reported that their cross-sectional profile may flatten above a certain threshold (black arrow head), thus softening the tube. **F)** Lattices of GDP-tubulin are 1-3% shorter than MTs that were polymerised with the non-hydrolysable GTP analogue GMPCPP, or stabilised with taxol (orange structure binding α-tubulin 1:1, according to Nogales et al., 1995); binding of kinesin-1 causes similar lengthening of tubulin (and additional compactions in the tubulin structure: yellow stars) which may cause cooperative binding of further kinesins and induce curvature if occurring only on one side of the MT; in extended taxol-bound MTs, bending forces were suggested to transfer tubulins on the concave side into their short conformation as an energetically favoured condition. For further references see main text.

Furthermore, MTs are structurally active: their physical properties can change when proteins bind to them (e.g. kinesins, see below) or when the ‘tubulin code’ is altered. The tubulin code is determined by the incorporation of different existing isotypes of α-and ß-tubulin into the MT lattice, and the addition of a range of distinct post-translational modifications (Fig.6B; Janke and Kneussel, 2010; Park and Roll-Mecak, 2018; Ti et al., 2018; Vemu et al., 2017). Some modifications influence the interaction with MT-binding proteins (e.g. poly-glutamylation attracts spastin; Valenstein and Roll-Mecak, 2016), others are believed to structurally protect MTs from damage or depolymerisation, such as poly-aminations on various residues (Song et al., 2013), or acetylation of luminal lysine 40 which has been suggested to make MTs more flexible and break-resistant (Fig.6B; Baas et al., 2016; Howes et al., 2014; Portran et al., 2017; Soppina et al., 2012; Xu et al., 2017). Notably, site-directed mutation of lysine 40 in *Drosophila* α1-tubulin could demonstrate that intraluminal MT acetylation is physiologically relevant (Jenkins et al., 2017; Yan et al., 2018). In addition, the MT lumen may contain MIPs (MT inner proteins) that likely also modify MT stability (Ichikawa and Bui, 2018).

These intrinsic or acquired physical properties are likely to determine how MTs respond to external forces - and we can expect such forces to be highly enriched in axons (see next section). Some ideas about how forces may impact on axonal MTs can be derived from *in vitro* experiments. For example, MTs in flow chambers that are anchored at one end, will bend when applying flow and rapidly return to straight confirmation thereafter; when bent repeatedly in this way, MTs experience structural damage that triggers subsequent repair responses (Akhmanova, 2018; Schaedel et al., 2015; Triclin et al., 2018); when certain shaft-binding proteins (e.g. doublecortin or non-motile kinesin-1) are added, MTs become locked in bent conformation and fail to re-straighten (Bechstedt et al., 2014; Ettinger et al., 2016; Peet et al., 2018).

Another example is provided by so-called *in vitro* gliding assays, where MTs are moved around on carpets of active motor proteins. On carpets of (axonemal) dynein, MTs move plus-end-first; they undergo collisions at high frequency, but seem to stay fairly straight and form vortices at the millimetre scale (Sumino et al., 2012). In contrast, if similarly prepared MTs are on kinesin carpets, they move minus-end-first and undergo fewer collisions because they can pass over one another, likely owed to kinesin-1’s adaptable length (Kerssemakers et al., 2006; Palacci et al., 2016; Sumino et al., 2012); however, if they collide or become pinned to the substrate (e.g. by dead kinesins) they frequently undergo dramatic shape changes at the micron-scale, including fishtailing and arc or loop formation (Amos and Amos, 1991; Lam et al., 2016; Weiss et al., 1991). The smallest diameters of curvature observed are similar to those of curled MTs in axons with values as low as 1-3 μm (Tab.1, Fig.5; Ahmad et al., 2006; Sánchez-Soriano et al., 2009) - and below 1μm, MTs are believed to break (Odde et al., 1999; Waterman-Storer and Salmon, 1997).

**Tab. 1.** MT loop or spool formation in gliding assays under different conditions. a) Primarily the lower range of mentioned diameters is listed; b) not clear from experimental section; c) measured from images. References [1] (Amos and Amos, 1991), [2] (Liu et al., 2011), [3] (Rashedul Kabir et al., 2012), [4](Kawamura et al., 2008), [5] (Hess et al., 2005), [6] (Lam et al., 2014), [7] (Wada et al., 2015), [8] (Luria et al., 2011), [9] (VanDelinder et al., 2016b), [10] (Liu et al., 2008), [11] (Liu and Bachand, 2013), [12] (Sumino et al., 2012). Note that a number of mathematical models were put forward to describe loop or spool dynamics in gliding assays (Crenshaw et al., 2011; Gosselin et al., 2016; Luria et al., 2011; Pearce et al., 2018; Ziebert et al., 2015).

| experimental conditions | diameters of curvature [$\mu\text{m}$]<sup>a</sup> | comments | ref. |
| --- | --- | --- | --- |
| **kinesin-1 carpets** |  |  |  |
| standard tub, 10-20 $\mu\text{m}$ taxol (after?)<sup>b</sup> polym. | 1-1.4<sup>c</sup> | waves and curls upon pinning | [1] |
| standard tub, 50 $\mu\text{M}$ taxol during & after polym.; high MT density (2.5 MTs/$\mu\text{m}^2$) | 1-5 | loops form through collision; loop duration up frequently >5 min; strong increase in loops at high MT conc.; decreasing loop radius with increasing contour | [2] |
| rhodamine-tub, 10 $\mu\text{m}$ taxol after polym.; exposing to air bubble or n-heptane | 1.1 (heptane), 1.8 (air) | MTs become reversibly unstable in non-polar conditions: 50% of MTs form loops as long as close to air bubble; effect absolutely requires kinesins | [3] |
| rhodamine-tub, 10 $\mu\text{M}$ taxol after polym. | 2.5-3.75<sup>c</sup> | left-handed supertwist favours CCW rotation of loops; CCW rotation is preserved in spools | [4] |
| biotin-tub, 10 $\mu\text{m}$ taxol after polym.; SA-linked | 1-12.6, mean 3.9 | | |
| biotin-tub, 10 $\mu\text{m}$ taxol after polym.; SA-linked | 1-5, mean 2.3 | up to 25 $\mu\text{m}$ long straight bundles; pinning of tip induces spools or fishtailing; rare "unspooling" events | [5] |
| biotin-tub, 10 $\mu\text{m}$ taxol after polym.; SA-linked; 1600, 870, 270 and 90 kinesins/$\mu\text{m}^2$ | ca. 2.4-4 | highest spool density & lowest spool diameter @ highest kinesin density; pinning as main cause for spool formation | [6] |
| biotin-GTP-tub, 10 $\mu\text{m}$ taxol after polym.; SA-linked | 5.7 (@ 10.8 $\mu\text{m}$ length), 3 (@ 3.7 $\mu\text{m}$) | spool diameters increase with MT length per condition; spool diameters: GMP-MTs (taxol) < GMPCPP-MTs (no taxol) < | [7] |
| biotin-GMPCPP-tub, 10 μm taxol after polym.; SA-linked | 18.8 (@ 10.3 μm length), 5.8 (@ 3.4 μm) | GMPCPP-MTs (taxol) | |
| biotin-GMPCPP-tub, no taxol; SA-linked | 8.2 (@ 10 μm length), 4.3 (@ 3.4 μm) | | |
| biotin-GTP-tub, 10 μm taxol (after?)<sup>b</sup> polym.; SA-linked | 3.2 μm (@ 6 μm length) | live imaging: pinning & collisions (simultaneous sticking) cause spool formation; spool formation is not activated by a Brownian ratchet type process | [8] |
| biotin-tub, 10 μm taxol after polym.; SA-linked; microfluidic device | 2.7 (pinning), 6.2 (collisions) | live imaging: pinning & collisions (simultaneous sticking) cause spools of different diameters; pinning more frequent in flow cells than microfluidic device | [9] |
| biotin-tub, (taxol?)<sup>b</sup> polym.; SA-quantum dot-linked | 1.2, mean 3.4 | left/right-handed super-twist: CCW/CW rotation; rings form intertwined wreath-like structures; tendency to disassemble involving MT breakage, kinesins pulling (blocked by AMP-PNP), counteracted by SA (enhanced by biotin) | [10] |
| biotin-tub, 10 μm taxol after polym.; SA-quantum dot-linked; patterned kinesin carpets | 1-5.3 and 3.1 | smallest spool diameters on constrained carpets: 1-5.3 μm on 5 μm stripes, 3.1 μm on 2 μm wide squares | [11] |
| **axonemal dynein carpet** |  |  |  |
| Cy3-tub, 10 μM taxol | straight | forming vortices in mm range | [12] |

If MTs on kinesin carpets are reversibly cross-linked with biotin-streptavidin, they coalesce into bundles containing dozens of MTs which frequently curl up into spools with inner diameters similar to those of loops. Spools can take on similar appearances as looped MT bundles observed in growth cones of fly or mammalian neurons (Dent and Kalil, 2001; Hess et al., 2005; Sánchez-Soriano et al., 2010). Furthermore, single MTs can escape from spools which may trigger spool disassembly (Hess et al., 2005; Liu et al., 2008; VanDelinder et al., 2016b), bearing some resemblance with off-track MTs in axons.

Loops and spools *in vitro* might therefore be an experimental proxy for curling MTs or bundled loops in axons, and gliding assays might provide mechanistic insights into these MT behaviours (details in Tab.1): for example, loop formation is favoured by high density of MTs and/or kinesins (Lam et al., 2014; Liu et al., 2011), and both are clearly given in axons; kinesins directly impact on MTs (see below), but they can also cause pinning events in gliding assays, which could be seen as a potential proxy for the abundant obstacles in the narrow axons. Furthermore, the diameters of curls and spools in gliding assays increase with the degree of MT rigidity (Wada et al., 2015), and their clockwise versus counter-clockwise directionality of circle formation is a function of the right-versus left-handed supertwist of the MTs involved - all properties that could potentially be tested in axons (Fig.6D; Kawamura et al., 2008; Liu et al., 2008). Furthermore, exposure to non-polar interfaces (e.g. n-heptane or air bubbles) induces strong curling (Rashedul Kabir et al., 2012), and this may be relevant in axons: in ageing or degenerative disease, changes in physical and chemical parameters of neurons affect liquid-liquid phase separation (Alberti and Hyman, 2016); liquid compartments likely are of low polarity (Nakashima et al., 2019) and might therefore influence the curling bias of MTs.

MT loops in gliding assays can be surprisingly stable (frequently >5 mins, as reported in Liu et al., 2011). To explain this, it has been proposed that tubulin-heterodimers on the concave side of the tube take on a shorter conformation than those on the convex site, and that this asymmetric distribution can be maintained as an energetically favoured state (Fig.6F, bottom right; Ziebert et al., 2015). In support of this model, tubulins in non-hydrolysed GMPCPP-MTs were shown to be 1-3% longer than hydrolysed GDP-tubulin, and taxol added after (but not during) polymerisation achieves a similar elongation (Fig.6F; Alushin et al., 2014; Amos and Löwe, 1999; Arnal and Wade, 1995; Castle et al., 2017; Hyman et al., 1995). Notably, this conformational length change seems physiologically relevant, as its suppression by the T238A mutation in yeast ß-tubulin stabilises MTs *in vivo* and causes mitotic defects (Geyer et al., 2015; Machin et al., 1995).

Such intrinsic properties of MTs may contribute to MT curling in axons, but we also need to consider the presence of MT lattice-associating proteins, such as tau, doublecortin or kinesin-1 which were reported to bind differently to straight and curved MTs (Balabanian et al., 2017; Bechstedt et al., 2014; Ettinger et al., 2016; Peet et al., 2018; Samsonov et al., 2004). In particular kinesins-1 was shown to stabilise MT curvature by extending their lattice to similar degrees as taxol (Peet et al., 2018), involving local compaction of tubulin that goes beyond taxol- or GMPCPP-induced effects (Krebs et al., 2004; Morikawa et al., 2015). Since kinesin-1 has a preference for convex MT surfaces and was reported to undergo cooperative binding, this may lead to a curvature-enhancing and -stabilising snowball effect with an estimated diameter of curvature of 3.2 μm (Cross, 2019; Muto et al., 2005; Peet et al., 2018). Mathematical modelling suggests that the kinesin carpet in gliding assays might induce stable yet reversible curling in this way (Fig.6F, top right; Pearce et al., 2018).

Naturally, current models are in their infancy and further findings need to be incorporated. For example, MTs behave as elastic cylinders (comparable to a garden hose) and can undergo softening through cross-sectional flattening when strongly bent (Fig.6E; Kononova et al., 2014; Memet et al., 2018). In this same vein, conformational changes of MTs upon kinesin-1 binding were reported to soften MTs locally (Kabir et al., 2014). If confirmed, this would have important implications for any existing models; together with the kinesin-induced tubulin compaction (yellow asterisks in Fig.6F), it might be a mechanism to absorb energy and reduce the shear force load on MTs. Notably, softening of MTs is also observed upon taxol application (usually used in gliding assays; Tab.1; Castle et al., 2017) or MT acetylation (abundant in axons; Portran et al., 2017; Xu et al., 2017), and might be a common prerequisite for curling behaviours.

Loop and spool formation in gliding assays are considered processes of ‘active self-organisation’ (Lam et al., 2016); given the above listed similarities, the same might be true for the formation of MT disorganisation in axons. Any *in vitro* studies addressing MT bending can provide potential mechanisms that could underlie MT curling in axons; gliding and flow chamber assays both suggest motor proteins, in particular kinesins, as key factors. In the next section we will therefore summarise roles of axonal MT-associated motors during axon pathology.

### The intricate relationship between MTs and their associated motor proteins

MT-associated motors comprise the minus end-directed dynein/Dynactin complex and the mostly plus-end directed proteins of the kinesin family (Hirokawa et al., 2010). Several kinesins display direct roles in MT regulation (Sturgill and Ohi, 2013). These include active MT depolymerisation (kinesin-8, −13; Walczak et al., 2013), MT polymerisation (kinesin-2, −5; Chen and Hancock, 2015; Gumy et al., 2013; Guzik-Lendrum et al., 2017), MT-cross-linkage (kinesin-5, −6, −12; see section on bundling), and roles in promoting MT orientation as a feature of neuronal polarity (Tas et al., 2017; Zheng et al., 2008).

However, most attention is given to the active cargo and organelle transport and dynamics in axons (Fig.3A-E; see section on axonal cytoskeleton), which is driven retrogradely by dynein/Dynactin (Allan, 2011) and anterogradely by kinesins (primarily kinesin-1, −2, and −3; Hirokawa et al., 2010). The forces imposed by these dynamics and/or the size of cargoes moved, poses an obvious challenge to MT bundles (Appert-Rolland et al., 2015) and might be an important factor leading to MT disorganisation.

Clearly, there is an intricate mutual regulatory relationship and finely tuned balance between the amount of transport, and the structural properties of the transport highways (Appert-Rolland et al., 2015; Prokop, 2013). On the one hand, MTs influence transport: for example, MT density is higher in small calibre axons than in large axons (∼15 *versus* ∼150 MTs/μm^2^), and mathematical modelling suggests that this is required to achieve the same transport efficiency as in large axons (Wortman et al., 2014; and references within); furthermore, the tubulin isotype composition of MTs, their posttranslational modifications, and the physical presence of other MT-binding proteins influence motor protein dynamics (’a’ in Fig.3; Balabanian et al., 2017; Monroy et al., 2018; Sirajuddin et al., 2014; Subramaniyan Parimalam et al., 2016). *Vice versa*, transport affects MT bundles: for example, kinesin binding changes the physical properties of MTs (see previous section), and motor proteins cause damage to the MTs they walk on, triggering maintenance responses including MT repair or potentially even replacement (’14’ in Fig.3; Akhmanova, 2018; Dumont et al., 2015; Peet et al., 2018; Triclin et al., 2018; VanDelinder et al., 2016a).

Tipping the balance in this mutual relationship can easily be imagined to cause reciprocal deficiencies in transport rate and MT bundle organisation. For example, disorganisation or partial breakage of MTs has been reported to cause pathological transport deficits (option ‘1’ in Fig.4; Fiala et al., 2007; Tang-Schomer et al., 2012). *Vice versa*, immunological lesioning experiments to induce demyelination (Abdul-Majid et al., 2000; Baker et al., 1990), initially caused transport defects, which were then followed by MT disorganisation (’2’ in Fig.4; Sorbara et al., 2014). Analogously, we observe severe MT disorganisation in *Drosophila* primary neurons upon loss of kinesin-1 or −3 (’2’ in Fig.4; unpublished; kinesin-1 shown in Fig.5E).

How loss of these kinesins may cause MT disorganisation can currently only be hypothesised. For example, it has been reported for dendrites that kinesin-1 migrates on acetylated and kinesin-3 on tyrosinated MTs (Tas et al., 2017). Provided the same is true in axons, the loss of kinesin-1 would relieve acetylated MTs, but tyrosinated MTs would still bear their full transport load - and *vice versa*. Such imbalances in transport distribution within MT bundles could lead to shear forces that buckle MTs and seed MT disorganisation. In the same vein, MT disorganisation was reported to be triggered by directional changes in motor traffic upon deficiency of the dynein regulator NDEL1 at the axon initial segment (Kuijpers et al., 2016). Furthermore, the movement of large cargoes likely induces dynamic and transient rearrangements of local MT-MT crosslinking networks (see section on cross-linkage) to make the necessary space; in this scenario, violating the balanced proportion between cross-linkers and transport may be a path to bundle aberration.

Alternatively, transport defects might affect MTs through biochemical routes, simply caused by the fact that the bundle-maintaining machinery runs out of supply and/or regulators. One would expect deficient supply (a) of tubulin heterodimers as building blocks, (b) of the proteins required to execute MT bundle maintenance work (’b’ in Fig.3), and an absence or wrong distribution (c) of organelles which are expected to play major roles in MT bundle maintenance (see Box 3 for details).

#### Box 3. The intricate relationship between MTs and axonal organelles

Mitochondria are the main source for ATP (Sheng, 2017), required to fuel multiple processes relevant for MT regulation (red arrows in Fig.3); these include actin assembly and dynamics relevant for MT regulation (Krendel and Mooseker, 2005; Skruber et al., 2018), protein phosphorylation for example of MT regulators (Bogoyevitch and Fairlie, 2007), GTP production required for MT polymerisation and signalling (Berg et al., 2002; Hall and Lalli, 2010; Voelzmann et al., 2016a), MT severing (McNally and Roll-Mecak, 2018), and MT-motor dynamics (Hirokawa et al., 2010; but note that vesicular transport uses local glycolysis to generate its own ATP; yellow star in Fig.3A; Hinckelmann et al., 2016; Zala et al., 2013). Secondly, the mitochondrial surface is an important signalling platform potentially required to orchestrate MT regulation locally (not shown in Fig.3; McBride et al., 2006). Thirdly, mitochondria cooperate with ER in the regulation of intracellular free calcium (yellow cloud in Fig.3; Rieusset, 2017; Wu et al., 2017) which has direct impact on MT regulators (e.g. spectraplakins, tau, kinesins; Kapur et al., 2012; McVicker et al., 2015) or even on MTs themselves (O’Brien et al., 1997). Fourthly, mitochondria collaborate with peroxisomes in the regulation of reactive oxygen species (’ROS’ in Fig.3; Fransen et al., 2017; Pascual-Ahuir et al., 2017), which have known effects on MT regulation (Wilson and Gonzalez-Billault, 2015). If excessive amounts of the wrong ROS species are produced upon transport-induced mitochondrial damage or dysregulation of the mitochondria-peroxisome system, this causes oxidative stress as a major path to axon pathology (Fiala et al., 2007; Liu et al., 2017; Pascual-Ahuir et al., 2017).

Such causative relationships between MTs and oxidative stress can be experimentally demonstrated: for example the MT-stabilising drug epothilone B rescues oxidative stress caused by peroxisome transport deficiencies in a human iPSC model of SPG4 (spastin-linked spastic paraplegia 4; Wali et al., 2016), suggesting that MTs are the cause for the transport deficit in the first place.

Similar interdependencies would apply to other important organelles or membrane compartments that likewise depend on MT-binding motor proteins to undergo meaningful dynamics (Fig.3D); of particular importance are the ER with its multiple roles in calcium homeostasis, protein synthesis and lipidogenesis (Gonzalez and Couve, 2014), or the endolysosomal system required for proteostasis (Winckler et al., 2018). Also drug-induced inhibition of the proteasome-ubiquitination system has been shown to induce alteration in MTs and axonal transport (Poruchynsky et al., 2008; Staff et al., 2013).

Functional interdependencies between transport, organelle dynamics and MTs provide potential explanations for a number of observations. For example, they may explain why axonal swellings induced by senile plaques in the *Tg-swAPP^Prp^* mouse (overexpressing an amyloid precursor protein carrying a familial Alzheimer’s disease-linked mutation; Stokin et al., 2008) were strongly enhanced when removing one copy of the KLC1 gene (a linker required for kinesin-1 mediated transport) - and this effect is conserved in *Drosophila* (Stokin et al., 2005). They may explain why different types of Charcot-Marie-Tooth disease or hereditary spastic paraplegias can be caused through motor proteins as well as regulators of membranous compartments (Blackstone, 2018; Bucci et al., 2012). They may also explain why MT stabilising drugs can be beneficial in animal models of neurodegeneration as diverse as SPG4 (Box 3) and Alzheimer’s disease (Brunden et al., 2014).

Naturally, the argumentative framework presented here is highly speculative, given the enormous complexity of the relationships between MT bundle organisation, motor protein activity and systemic factors. But we hope that these reflections will motivate experimenters to have a closer look at MTs in future studies of axon biology and pathology, and include statements in their reports as to whether MTs are affected. More data are urgently needed, which does often not require more than analysing neuronal morphology with antisera against MTs (rather than restricting to intermediate filaments), or increasing the magnification in ultrastructural studies to have a closer look at MTs. In the following sections we will explore the mechanisms that are potentially used to form and maintain MT bundles against the odds of motor-induced aberration or damage.

### MT polymerisation as a fundamental requirement for bundle maintenance

The *de novo* formation of MT bundles during developmental, plastic or regenerative axon growth (’8’ in Fig.3) requires MT polymerisation. At later stages, MTs continue to undergo polymerisation {Kleele et al., 2014; Voelzmann et al., 2016a}, likely to maintain a steady state and prevent MT senescence through polymerisation-dependent MT repair and/or turn-over (’14’ in Fig.1; Akhmanova, 2018; Triclin et al., 2018). A well-regulated machinery of MT polymerisation and disassembly (blue stippled arrows in Fig.3) is therefore needed to keep the numbers of axonal MTs in balance with the transport load (see previous section; Wortman et al., 2014).

As we detailed in a previous review (Voelzmann et al., 2016a), the machinery of MT de-/polymerisation requires three sub-machineries: (1) dynamic protein complexes at the MT plus end (blue balls, ‘Eb1’ in Fig.3); (2) the supply of α/β-tubulin heterodimers as building blocks which occurs through a complex regulatory network in close co-regulation with MT dynamics (’c’ in Fig.3; Al-Bassam, 2017; Gasic and Mitchison, 2018; Preitner et al., 2014); (3) proteins which bind or post-translationally modify the MT lattice with impact on plus end dynamics, for example by stabilising MTs against depolymerisation (‘7’ in ‘Fig.3).

The fine-tuning of net MT polymerisation appears to depend on complex regulatory networks. This is illustrated by our recent work in *Drosophila* neurons, showing that loss of cortical actin rings in the axon shaft (Fig.1) causes a reduction in MT polymerisation speed, eventually affecting MT bundle integrity; simultaneous application of MT-destabilising drugs or removal of the MT-stabilising spectraplakin Short stop (Shot) exacerbated these effects, frequently even eliminating entire axons (Qu et al., 2017). Similar dependencies of MT polymerisation on actin networks are suggested by other reports: (1) parallel loss of spectrin and tau causes axonal MT loss in *C. elegans* (Krieg et al., 2017); (2) axon-shortening induced by the MT-stabiliser taxol can be ameliorated through co-application of actin-destabilising drugs (in both chick and *Drosophila* neurons; Letourneau et al., 1987; Sánchez-Soriano et al., 2010); (3) application of actin-destabilising drugs changes the tubulin-to-microtubule ratio in PC12 cells (Dennerll et al., 1988) and causes axon retraction in chick dorsal root ganglia neurons (Datar et al., 2019; see also Box 4). The best explanations for the mechanistic links from actin networks to net MT polymerisation are currently provided by biomechanical models (see Box 4).

#### Box 4. Biomechanical models of axon growth

The regulation of axonal growth dynamics has been explained in terms of balance of forces between the microtubule and actin cytoskeleton (de Rooij et al., 2018; Fan et al., 2017; Miller and Suter, 2018). In axons, “*actin is under tension supported in part by microtubules under compression*” (Dennerll et al., 1988; Heidemann and Buxbaum, 1990). Tension is provided by the pull of the growth cone (Chan and Odde, 2008; Koch et al., 2012; Lamoureux et al., 1989) and the active contraction of acto-myosin along the axon shaft (Fan et al., 2017; O’Toole et al., 2015; Fig.1); the stiff nature of cross-linked MT bundles is well suited to oppose compressive forces up to a certain threshold (Buxbaum and Heidemann, 1992; de Rooij et al., 2018; Fig.2).

In such a balanced system, manipulations such as externally imposed pulling forces (Bray, 1984; Lamoureux et al., 2010; Pfister et al., 2004; Zheng et al., 1991) or genetic/pharmacological de-/stabilisation of acto-myosin (Ahmad et al., 2000; Datar et al., 2019; Dennerll et al., 1988; Ketschek et al., 2007; Turney et al., 2016; Wylie and Chantler, 2008) clearly modulate axon length or growth. Part of this response is expected to be due to changes in MT assembly, as was found when applying external forces to non-neuronal cells (Kaverina et al., 2002). But MTs themselves are not only responders in this context: the dis-/assembly or motor-based sliding of MTs can actively contribute by generating forces (Ahmad et al., 2000; Brouhard and Rice, 2018; Roossien et al., 2014; Winding et al., 2016).

How forces are sensed and translated into compensatory force generation and/or changes in axonal length or growth, remains an important question (see also the last section on cortical anchorage). Potential mechanisms might involve mechanically induced conformational changes of MTs (single MTs polymerise faster when being pulled on *in vitro*) or responses of polymerases such as XMap215 (Brouhard and Rice, 2018); but especially mechano-sensitive calcium channels in the axonal membrane (Franze et al., 2009; He et al., 2019; Heidemann and Buxbaum, 1990; Song et al., 2019) are promising candidates to orchestrate local responses that can even go beyond mere changes in MT polymerisation.

### Maintaining MTs bundles through cortical guidance and elimination of polymerising MTs

Whilst MT polymerisation is a requirement for axon formation and maintenance, it also poses a risk: for example, extending MTs may be obstructed by the abundant organelles or protein complexes in axons, thus causing accidental ‘off-track’ MTs that project out of the bundle towards the cortex (’4’ in Fig.3). Apart from MT buckling, off-track MTs may be a second cause for axonal MT disorganisation.

A key factor preventing this from happening is Eb1 (Alves-Silva et al., 2012; Figs.3 and 5B). Eb1 directly binds at extending MT plus ends where it promotes polymerisation (Zanic et al., 2013) and serves as a scaffold for many other proteins (Gupta et al., 2014). In the absence of Eb1, MTs are severely disorganised, indicating important roles in MT maintenance (Alves-Silva et al., 2012). One underlying mechanism is the guidance of polymerising MTs through binding of Eb1 to Short stop (Shot); Shot is a well-conserved spectraplakin, able to cross-link cortical actin, MTs and Eb1 (’5’ in Fig.3), thus guiding polymerising MTs in parallel to the axonal surface and laying them out into parallel bundles (Alves-Silva et al., 2012). Accordingly, also loss of Shot causes severe MT disorganisation in axons - and the same is true for its two mammalian homologues ACF7 and dystonin (Bernier and Kothary, 1998; Dalpe et al., 1998; Sánchez-Soriano et al., 2009; Voelzmann et al., 2017) - of which the latter links to the axonopathy HSAN6 (type 6 hereditary sensory and autonomic neuropathy; Edvardson et al., 2012).

Such cortical guidance is complemented by at least one control mechanism: if MTs (accidentally) leave their bundled arrangements and extend towards the cortex, they get eliminated by Efa6 (’4’ in Fig.3), a cortical collapse factor that associates with the axonal membrane via its C-terminal plekstrin homology domain; when Efa6 is absent, off-track MTs outside axonal MT bundles persist for longer and are higher in number (Qu et al., 2018). Consistent with the known roles of off-track MTs in axon growth, branching and MT disorganisation (see Box 1 and above), neurons lacking Efa6 display longer axons, more branches and prominent MT disorganisation (Fig.5D; Qu et al., 2018).

Our model would predict that mutant phenotypes caused by loss of Shot and Efa6 should enhance each other because they are caused through complementary mechanisms of MT bundle regulation. Accordingly, we found a clear increase in MT disorganisation when removing both Shot and Efa6 from the same neurons, in culture and *in vivo* (Qu et al., 2018). We propose therefore that Shot and Eb1 keep MTs away from the membrane, whereas Efa6 acts as a quality control factor eliminating occasional accidental off-track MTs; this elimination seems to occur in moderate, well-balanced amounts so that ‘intended’ off-track MTs required for axon growth and branching can persist and perform their function.

Interestingly, the cortical collapse function of Efa6 is not conserved in vertebrates (Qu et al., 2018). Nevertheless, the concepts derived from Efa6 studies appear relevant, because loss of the unrelated neuronal cortical collapse factor KIF21A (a type 4 kinesin) causes analogous phenotypes in mammalian neurons. Thus, KIF21A mutations linked to the neurodevelopmental disorder CFEOM1 (type 1 congenital fibrosis of the extraocular muscles) affect axon growth and axonal branching just like Efa6 (Qu et al., 2018; van der Vaart et al., 2013) - and might as well cause MT disorganisation, but no data are currently available.

However, guidance along cortical actin seems not the only mechanism through which Eb1 and Shot keep MTs on track. This is illustrated by the simple fact that MT disorganisation observed upon loss of Shot or Eb1 (Fig.5B, E) does not occur when removing actin from axon shafts (Alves-Silva et al., 2012; Qu et al., 2017; Sánchez-Soriano et al., 2010). This suggests that both factors perform additional, actin-independent functions or interactions to promote MT bundles.

For example, the unusual Shot-PH isoform, which is highly enriched in the nervous system and harbours a plakin repeat region (PRR; conserved in mammalian spectraplakins), is a likely candidate for such roles that still await investigation (’11’ in Fig.3; Hahn et al., 2016; Voelzmann et al., 2017). Eb1 has a long list of protein interactors besides Shot (Gupta et al., 2014), and some of them might associate with MTs and guide extending plus ends along pre-existing bundles (’9’ in Fig.3); for example, APC or GAS2-LIKE family members (Pickled eggs/Pigs in *Drosophila*) are good candidates, known to bind both MTs and Eb1 in mammals and *Drosophila* (Beaven et al., 2015; Pines et al., 2010; Stroud et al., 2014).

### Potential roles of severing proteins and MT-destabilising kinesins in MT bundle maintenance

Apart from cortical MT elimination, also MT severing and/or depolymerisation in the cytoplasm may play important roles in maintaining axonal MT bundles. This is supported by axonal MT disorganisation observed upon the losses of *Drosophila* katanin (our unpublished results) or mammalian spastin (Denton et al., 2014; Fassier et al., 2013; Havlicek et al., 2014; Tarrade et al., 2006).

As explained in the previous section, MTs leaving the bundled conformation can drive axonal growth, branching and MT disorganisation, and cortical collapse factors negatively regulate all three processes. In line with this argumentation, also the MT-depolymerising kinesin-13 family member Kif2A (Homma et al., 2003) and MT severing proteins (spastin, katanin and fidgetin) were reported to inhibit neurite growth and/or branching (Leo et al., 2015; Mao et al., 2014; Tao et al., 2016). However, other studies of spastin, katanin and fidgetin led to contradictory findings, describing them as promoters rather than inhibitors of neurite growth and branching (Ahmad et al., 1999; Butler et al., 2010; Havlicek et al., 2014; Karabay et al., 2004; Riano et al., 2009; Stewart et al., 2012; Stone et al., 2012; Wood et al., 2006; Yu et al., 2008). Such stark, potentially context-dependent deviations seem to reflect the complex regulation of severing proteins.

Spastin, katanin and fidgetin are all members of the superfamily of AAA proteins (ATPases associated with diverse cellular activities; McNally and Roll-Mecak, 2018; Sharp and Ross, 2012; Zhang et al., 2007), but their severing activity is differentially regulated through their individual responses to (a) posttranslational MT modifications (in particular acetylation and poly-glutamylation; Bailey et al., 2015; Lacroix et al., 2010; Leo et al., 2015; Shin et al., 2019; Sudo and Baas, 2010; Valenstein and Roll-Mecak, 2016), (b) antagonistic MT shaft-binding proteins such as tau (Qiang et al., 2018; Qiang et al., 2006; Yu et al., 2008), or (c) spatial recruitment through specifically localised proteins such as CAMSAP (Jiang et al., 2018). Furthermore, katanin has the ability to depolymerise MTs in an ATP-independent manner (Belonogov et al., 2019).

Through this precise context-dependent spatiotemporal regulation of their activities, severing proteins can have two diametrically opposed outcomes: they either eliminate MTs and reduce their numbers, or they break them up into stable fragments that serve as seeds for MT amplification (Baas et al., 2016; McNally and Roll-Mecak, 2018; Vemu et al., 2018). In the following, we will briefly discuss how either of these outcomes could be used to prevent MT disorganisation:

First, MT severing proteins could complement roles of cortical collapse factors (’4’ in Fig.3) by serving as quality control factors that eliminate disorganised MTs in the cytoplasm (’6’ in Fig.3). For example, katanin in plant cells was reported to localise and sever preferentially at MT cross-points, which can be used to take out non-aligned MTs (McNally and Roll-Mecak, 2018).

Second, MT shortening functions of katanin are required at MT minus ends. Thus, in both mammals and *Drosophila*, the minus-end capper CAMSAP/Patronin protects against MT disassembly, and recruits katanin to counterbalance against uncontrolled minus-end polymerisation (’13’ in Fig.3; Goodwin and Vale, 2010; Jiang et al., 2018; Nashchekin et al., 2016); uncontrolled minus end extension upon katanin deficiency may cause MTs to go off-track or to buckle through extra forces produced.

Third, MT elimination functions could prevent MT bundle senescence. For example, MTs suffer from damage through tear-and-wear (Dumont et al., 2015; Peet et al., 2018; Schaedel et al., 2015; Triclin et al., 2018; VanDelinder et al., 2016a), which might cause bundle aberration by abrogating interactions with MT-binding proteins (red cross at ‘16’ in Fig.3). MT fractures or holes can be repaired through mechanisms involving katanin or spastin (Davis et al., 2002; Diaz-Valencia et al., 2011; Gasic and Mitchison, 2018; Triclin et al., 2018; Vemu et al., 2018). More subtle features of senescence (e.g. irreversible modifications, loss of tubulin C-tails) might require selective elimination of ageing MTs through severing factors (as similarly suggested for kinesin-8 or −13; Gardner et al., 2011), followed by compensatory polymerisation (’14’ in Fig.3). For example, spastin deficiency in the *Sp^Δ^* mouse model causes a drop in MT polymerisation (which might reflect reduced turn-over) accompanied by a rise in MT disorganisation (which might be due to precocious MT senescence; Fassier et al., 2013).

However, the MT phenotypes observed in the *Sp^Δ^* mouse model could likewise be explained through the opposite role of spastin in MT multiplication. Thus, in the absence of spastin-mediated amplification, MT numbers might gradually decline and cause transport interruptions; this, in turn, would affect MT bundle organisation and eventually cause axonal pathology (see section on motor proteins; Wali et al., 2018; Wali et al., 2016). Curiously, axon swellings in this model were reduced with low doses of MT-stabilising or -destabilising drugs (Fassier et al., 2013), therefore failing to provide any clues as to whether spastin works through MT turn-over or amplification in this context.

Understanding spastin is important because it is by far the most prominent factor linking to spastic paraplegias worldwide (Koh et al., 2018; Schüle et al., 2016), and axonal swellings are a hallmark of the disease (Blackstone, 2018; Zempel and Mandelkow, 2015). Most SPG4-linked mutations lie within the AAA-ATPase domain (Shoukier et al., 2009), suggesting that MT severing is key to the disease pathology. However, point mutations might generate versions of spastin, which either act as dominant negative alleles (forming dysfunctional complexes that titrate out other spastin-interacting factors), or acquire gain-of-function qualities by diffusing away to perform very different roles. One such MT-independent role of spastin is the isoform-specific regulation of the endoplasmic reticulum (’15’ in Fig.3), including its shape, its interaction with the endosome and its production of lipid droplets (Allison et al., 2017; Papadopoulos et al., 2015; Park et al., 2010; Solowska and Baas, 2015). It is therefore difficult to exclude that at least part of those SPG4-linked mutations triggers axon decay through other routes than the direct induction of MT aberrations (’2’ or ‘3’ versus ‘1’ in Fig.4).

### Potential roles of MT-MT cross-linkage in MT bundle maintenance

MT-MT cross-linkage (’12’ in Fig.3) is likely the oldest mechanistic concept put forward by neurobiologists to explain MT bundles (Lee and Brandt, 1992) and appears an obvious means of suppressing MT disorganisation. Physical cross-linking strands between axonal MTs were observed decades ago (Hirokawa, 1982; Hirokawa, 1986), and mathematical models support MT-MT cross-linkage as an important structural feature of axons (e.g. de Rooij and Kuhl, 2018; Lazarus et al., 2015; Li et al., 2018; Peter and Mofrad, 2012). To illustrate this point, axons have been described as a “*stiff spring in series with a viscoelastic (Voight) element composed of a less stiff spring in parallel with a fluid dashpot*” (Heidemann et al., 1990), meaning that axons are under rest tension and combine elastic and viscous properties. A central structural component underpinning such properties is likely provided by networks of MT-MT cross-linkers (Fig.2), where each linker is able to detach upon super-threshold pull or compression, and re-attach thereafter (slip-bonds). However, the molecular players mediating MT-MT cross-linkage remain surprisingly controversial to this day, as explained in the following.

First, showing that a neuronal linker expressed in non-neuronal cells induces MT bundling, is insufficient proof: MT bundling can even be achieved through expression of isolated MT-binding domains, or the application of the MT-stabilising drug taxol which causes bundles with ultrastructural cross-bridges that are indistinguishable from those induced by tau or MAP2 (Chapin et al., 1991; DeBonis et al., 2015; Goriounov et al., 2003; Kader et al., 2017; Lee and Brandt, 1992).

Second, dynamin is linked to Charcot-Marie-Tooth disease and has been shown to bundle MTs *in vitro*; however, the physiological relevance of this is questionable, because dynamin *in vivo* seems to bind primarily membranes (Scaife and Margolis, 1990; Shpetner and Vallee, 1989; Züchner et al., 2005).

Third, MTLC1 and MAP1B (Futsch in *Drosophila*) appear ideal cross-linkers, because they both possess an N-and a C-terminal MT-binding domain; they were shown to induce MT bundles upon expression in non-neuronal cells (with MAP1B being a weak bundler; Kader et al., 2017; Penazzi et al., 2016; Satake et al., 2017), and the fly homologue Futsch promotes looped MT bundles at synaptic terminals (Roos et al., 2000). Upon loss-of-function, MTLC1 causes MT disorganisation at the axon initial segment, strongly supporting its role as MT-MT cross-linker in this specific compartment (Satake et al., 2017). In contrast, the long history of MAP1B/Futsch research is mostly dedicated to aspects of axon development (Hummel et al., 2000; Migh et al., 2018; Villarroel-Campos and Gonzalez-Billault, 2014), but we are aware of only one isolated report showing axonal bundle defects (upon loss of Futsch; Bettencourt da Cruz et al., 2005).

Fourth, the other conserved linker candidate tau, has only one central MT-binding region, but it achieves physical MT-MT linkage *in vitro* through N-terminal dimerisation (Chung et al., 2016; Méphon-Gaspard et al., 2016; Rosenberg et al., 2008). However, its dwell time on MTs seems very short (Janning et al., 2014; Samsonov et al., 2004); similar to MAP1B/Futsch, reported tau-deficient phenotypes in neurons mainly concern developmental defects (Penazzi et al., 2016), but we are aware of only one report of bundle aberration (in *C. elegans*; Krieg et al., 2017; Penazzi et al., 2016).

Pinpointing roles of tau or MAP1B/Futsch in MT-MT cross-linkage is enormously complicated by the fact that both proteins seem to perform a whole array of further molecular functions relevant for MT dynamics. For example, tau can protect MTs from severing by katanin (Qiang et al., 2006), bind tubulin hetero-dimers (Shin et al., 2018), switch between bundled and single MT states (Prezel et al., 2018), cross-link MTs with actin or the cortex (Biswas and Kalil, 2018; Cabrales Fontela et al., 2017; Maas et al., 2000), stabilise MTs during axon initiation (Brandt, 1998), maintain labile domains along MT shafts (Qiang et al., 2018), regulate End-binding proteins (Sayas et al., 2015), compete with kinesins (Trinczek et al., 1999), and promote MT nucleation and polymerisation (Penazzi et al., 2016). A similarly broad pleiotropy has been reported for MAP1B (Villarroel-Campos and Gonzalez-Billault, 2014).

Pinpointing relevant MT-MT cross-linking activities of specific factors is also complicated by functional redundancies. For example, *MAP1B* and *tau* mutations have enhanced growth phenotypes when combined in double-mutant mouse neurons (Takei et al., 2000), and co-expression of Futsch and Tau causes enhanced phenotypes in the *Drosophila* CNS (Hummel et al., 2000). Functional redundancies likely extend to further potential cross-linkers. For example, Kinesin-5 (KIF11), kinesin-6 (KIF23, Pavarotti in *Drosophila*) and kinesin-12 (KIF15) slide anti-parallel MTs in the mitotic spindle (Baas, 1999); in axons MTs are arranged in parallel, and these kinesins seem to inhibit sliding (Dong et al., 2019; Lin et al., 2012; Liu et al., 2010; Lu et al., 2013; Myers and Baas, 2007; Nadar et al., 2012), indicating that they cross-link MTs. In support of this idea, we observe that loss of *Drosophila* Pavarotti causes axonal MT disorganisation which might reflect potential linker function (unpublished data).

In conclusion, MT-MT cross-linkage is a widely accepted concept, but experimental support for its existence in axons and our knowledge of the molecular players involved is insufficient. We even cannot fully exclude a model where MT bundles are held together by the corset of contractile cortical actin rings (Fig.1), and cross-linkers merely separate MTs to generate space for transport (Fan et al., 2017). If we are to decipher the true molecular nature of MT-MT cross-linkage in axons, future studies will have to address the challenges of functional redundancies between different classes of linker candidates.

### Does MT bundle maintenance involve their anchorage to the axonal surface?

Apart from cross-linking MTs within axonal bundles, they might also be anchored to the axon wall, thus achieving an even more stable structure that can prevent MT buckling and bundle deformation caused by the enormous forces imposed by axonal cargo transport. Relevant in this context are observations in developing vertebrate and fly neurons of a gradual flow of MT bundles towards the distal axon tip (Miller and Sheetz, 2006; O’Toole et al., 2008; Reinsch et al., 1991; Roossien et al., 2013). Forces contributing to this process could be derived from an increase in MT volume through polymerisation along the axon shaft (Sánchez-Soriano et al., 2010), pulling forces in the rear of growth cones (O’Toole et al., 2015), thermal motion of MT-MT cross-linkers (Lansky et al., 2015), kinesins actively sliding MTs along other MTs (’B’ in Fig.3; Lu and Gelfand, 2017), or dyneins sliding MTs along cortical F-actin (’10’ in Fig.3; Ahmad et al., 2006; He et al., 2005; Myers et al., 2006; Roossien et al., 2014).

Potential MT sliding along cortical actin would represent one form of tethering MT bundles to the axonal surface. Such anchorage is also suggested by observed co-drift of the axolemma with the axon core (Lamoureux et al., 2010; Popov et al., 1993; Zheng et al., 1991). But anchorage would not have to be static; for example, it might involve an interface of slip-bonds, as similarly suggested for actin networks that flow across, whilst dynamically anchoring to, relatively stable focal adhesion sites (Case and Waterman, 2015). MTs could anchor to cortical actin (Fig.1; ‘2’ in Fig.3; Xu, 2013 #6895} or to membrane-associated or transmembrane proteins including ion channels, ion transporters or adhesion factors (Fig.1; ‘3’ in Fig.3). Links to transmembrane proteins could be used as mechano-sensing modules (Yap et al., 2018) to measure local shear forces generated between MT bundles and the axonal environment (Fig.1). Such mechano-sensing could explain local regulation phenomena: for example, net rates of mitochondrial movement along the axon are fairly constant, but the slow transport component (driven by MT bundle flow) is low in proximal and high in distal axon segments; this gradual increase in the amount of slow transport is compensated for by fast transport (high proximal, low distal; Miller and Sheetz, 2006). The regional amount of fast mitochondrial transport could potentially be regulated by mechano-sensing, measuring the local MT drift rate relative to the outer axonal environment.

Apart from dynein (see above), other potential anchoring mechanisms can be deduced from the literature. For example, spectraplakins are good candidates, as suggested by distal shift of axonal MTs in fly neurons lacking the *Drosophila* spectraplakin Shot and treated with the MT-stabilising drug taxol (Voelzmann et al., 2017). Three distinct mechanisms could account for spectraplakin-mediated MT anchorage: Firstly, spectraplakins could directly cross-link actin and MTs (’2’ and ‘5’ in Fig.3). Secondly, they could link to membrane-associated proteins; thus, the mammalian spectraplakin dystonin can link to ß4-integrin and transmembrane collagen XVII (’3’ in Fig.3; Voelzmann et al., 2017), and *Drosophila* Shot is able to regulate the axonal localisation of the cell adhesion molecule Fasciclin 2, potentially cross-linking Fasciclin 2 to MT bundles (Bottenberg et al., 2009; Prokop et al., 1998). Thirdly, spectraplakins were shown in non-neuronal cells of fly and mammals to anchor MT minus ends to the cortex (’1’ in Fig.3; Nashchekin et al., 2016; Ning et al., 2016; Noordstra et al., 2016); this mechanism requires interaction with the MT minus end-stabilising factor CAMSAP/Patronin, a factor that is known to be relevant for neuronal morphology (Yau et al., 2014).

Also other MT-binding proteins, such as tau, MAP1B, APC and dynamin, might be involved in anchorage since they were also reported to bind to actin or to the cortex (’2’ in Fig.3; Biswas and Kalil, 2018; Blanchoin and Michelot, 2012; Brandt et al., 1995; Elie et al., 2015; Gu et al., 2010; Maas et al., 2000; Mohan and John, 2015; Villarroel-Campos and Gonzalez-Billault, 2014). But potential MT-actin cross-linkage in the axon may not only occur at the cortex, but as well at central longitudinal actin trails (Fig.1; Leterrier et al., 2017), thus further contributing to the intricate cross-linking networks expected to stabilise MT bundles. Deciphering MT bundle cross-linkage, internally or with the axonal surface, stays a major challenge for future experimentation that will teach us important lessons about axon biology and pathology.

### Conclusions and future perspectives

Here we have presented a conceptual view by describing a functional interactome that integrates the enormous complexity of cross-regulatory networks acting at the local level in axons. We propose that there has to be a fine balance between damaging effects inflicted by life-sustaining motor movements (’associated’, ‘A-E’ in Fig.3) and the activities that maintain the highways required for this movement (MT-’taming’ mechanisms; ‘1-15’ in Fig.3); both are fine-tuned through a number of cross-regulatory mechanisms (’a-e’ in Fig.3).

Our model integrates a broad range of findings from the literature, but its original foundations are derived from our own work in *Drosophila* neurons as one consistent cellular system. Like other genetic invertebrate models, *Drosophila* provides a cost-effective and fast system to unravel the functional overlap and interface of different genetic factors - ideal to dissect complex machinery and deliver data that then often apply to axons of higher animals (Beaven et al., 2015; Prokop, 2018; Prokop et al., 2013).

This strategy offers one feasible strategy towards solving the daunting task of disentangling the enormous complexity of axonal MT bundle regulation. For this, the model of local axon homeostasis could provide a useful basis, helping to develop testable working hypotheses; a good starting point might be to break down the local axon homeostasis machinery into classifiable sub-machineries, like those discussed in the different sections of this review.

This approach also means that the discovery of new molecular mechanisms should no longer be the only gold standard for axon research, but we need to recognise the value of long-term approaches that gradually assemble known and newly discovered molecular mechanisms into an integrated understanding of how regulation at the cellular level can be orchestrated. In our opinion, this would be a much needed strategy shift, providing understanding of axons at the organisational level at which axonopathies become manifest. As B.A. Cohen put it: “Research that results in models that reliably and quantitatively predict the outcomes of genetic, biochemical, or pharmacological perturbations should be valued highly, and rewarded, regardless of whether such models invoke novel phenomena” (Cohen, 2017).

For the studies of MTs in neurons, we need to take into consideration that knowledge derived from non-neuronal cells might not apply (Beaven et al., 2015). Furthermore, the interactome shown in Fig.3 makes clear that we will need quantitative approaches: we know increasingly well how factors bind to MTs and partly understand how they might compete with each other. But how crowded can a single MT be, how many molecules are there in its surrounding at any time point, and how much dynamic exchange is taking place? Computational modelling will be an unavoidable means to make sense of existing data and make reasonable predictions to inform experimentation (Cohen, 2004; Gunawardena, 2014).

Integrated understanding of axon biology will also improve our knowledge of the next higher level of complexity, i.e. the mechanisms that orchestrate axon homeostasis and that maintain balance even during phases of change (e.g. when switching from growth to differentiation, or during stress, injury, regeneration) - or that tip the balance and induce degeneration. Obviously, signalling networks or dynamic changes of systemic factors such as second messengers or the ‘tubulin code’ will be key players to this end (Baas et al., 2016; Park and Roll-Mecak, 2018; Schelski and Bradke, 2017; Wilson and Gonzalez-Billault, 2015) - and glial cells will likely act as important external influencers of such processes (Pan and Chan, 2017).

Finally, MTs have been recognised as promising therapeutic targets (Baas and Ahmad, 2013; Eira et al., 2016; Zempel and Mandelkow, 2015), and urgently needed advance on this translational path will be facilitated by a better understanding of the axonal MT homeostasis system. A larger focus of the research community on MTs, and generation of more and relevant data that can be incorporated into our understanding, would be a key prerequisite to make such progress.

## Acknowledgements

Work underpinning this article was made possible through support by the BBSRC to A.P (BB/I002448/1, BB/P020151/1, BB/L000717/1, BB/M007553/1), by parents to Y.-T.L., by the Leverhulme Trust to I.H. (ECF-2017-247). The Manchester Bioimaging Facility microscopes used in this study were purchased with grants from the BBSRC, The Wellcome Trust and The University of Manchester Strategic Fund. The Fly Facility has been supported by funds from The University of Manchester and the Wellcome Trust (087742/Z/08/Z). We thank Anthony Brown for introducing A.P. to the arguments that support a role of intermediate filaments in axon diameter, Anna Akhmanova for constructive and helpful feedback that has hopefully been well addressed in our last version, and Kyle Miller for major contributions to the contents of Box 4.

